# An high dose of a conjugated linoleic acid increases fatty liver and insulin resistance in lactating mice

**DOI:** 10.1101/588962

**Authors:** Kun Pang, Song bo Zhu, Liqiang Han

## Abstract

This study aimed to evaluate the effect of a high dose of conjugated linoleic acid (CLA) on lactating mice. In experiment one, KUNMING mice were separated into four groups (n = 6 per group); the control (CON) group received 3.0% linoleic acid oil (LA), the L-CLA group received 1.0% LA and 2.0% CLA mixture, the M-CLA group received 2.0% LA and 1.0% CLA mixture, and the H-CLA group received 3.0% CLA mixture. Feeding proceeded from day 4 to day 10 during lactation. In experiment two, the CON group received 2.0% LA and the H-CLA group received 2.0% CLA. Blood parameters were analysed for all groups, and insulin tolerance tests (ITTs) were conducted. CLA treatment did not affect the dam weight, but it significantly decreased the food intake of dams. Furthermore, CLA decreased the weight of pups, which was attributed to lower milk fat. H-CLA group mice displayed increased liver weight and liver triglyceride content, as well as a higher TG content and γ-GT activity in blood. Moreover, a high dose of CLA resulted in insulin resistance, possibly affecting the RBC and HCB of blood. In conclusion, lactating mice receiving a high dose of CLA led to fatty liver, insulin resistance, and impaired lactation performance.

## Introduction

Conjugated linoleic acids (CLAs) are molecules mostly found in meat and dairy products. CLAs have emerged as a possible adjuvant treatment for obesity because some studies show a reduction in body fat and increased lean mass after supplementation of a mixture of two isomers in the diet [1,2]. CLAs are also reported to affect carcinogenesis, glucose and lipid metabolism, diabetes, body composition, and immune cell functions [3–5]. However, certain isomers seem to cause fat accumulation in the enlarged liver of mice [6], and can induce insulin resistance in mice and humans [7,8].

In humans and other animals, milk lipids are markedly affected by the lipid composition of the diet, the maternal energy balance, and metabolism. Maternal lipid intake during lactation can alter milk composition, which in turn can affect the growth of the litter [9] and change lipid metabolism and liver enzymes [10]. Many previous studies showed that CLAs can lower triacylglycerol concentrations and alter the fatty acid profile in the milk of lactating animals including human, mouse, rat, sheep, goat and cow [11–17]. Feeding lactating mice with CLA can result in diminished mammary gland lipogenesis and reduced growth rate [18,19], but the exact metabolic and physiological changes involved remain unclear. More recently, Bezan (2018) demonstrated that feeding a high dose (3%) of CLA increases insulin resistance and exacerbates hepatic steatosis in male Wistar rats [20]. The intake of trans-fatty acids during lactation was also found to increase maternal lipids in the liver (Assumpção, 2004) and liver weight [21]. As indicated above, it is most likely that CLA not only affects mammary glands during lactation, but also liver and glucose metabolism.

CLAs used in supplements originate from the processing of vegetable oils such as Safflower oil, and always include an equimolar mixture of c9,t11-CLA and t10,c-12-CLA. Most studies of CLAs have used a mixture of isomers, with c9,t11-CLA and t10,c-12-CLA the most abundant [20,22]. To explore this further, two experiments were performed in which mice were fed with a CLA mixture. Firstly, the effects of different doses of the CLA mixture on the milk and liver of dams was investigated. Secondly, we tested whether a high dose of CLA could cause insulin resistance in dams.

## Materials and Methods

### Diets and feeding

Kunming (KM) mice were approved for use in this study by the University of Zhengzhou for Medical Sciences Animal Care and Use Committee. Animals were housed at 24°C with 40% humidity and a 12/12 h light-dark cycle. Animals were provided with food and water random. A mouse-breeder diet (Laboratory Animal Central, Henan province, China) containing the required complement of nutrients for lactation was used as a carrier for each of the four fatty acid treatments. The fatty acid composition of CLA oil and LA oil is shown in Table 1. Diets were prepared the day before parturition and stored at 4°C.

**Table 1.**
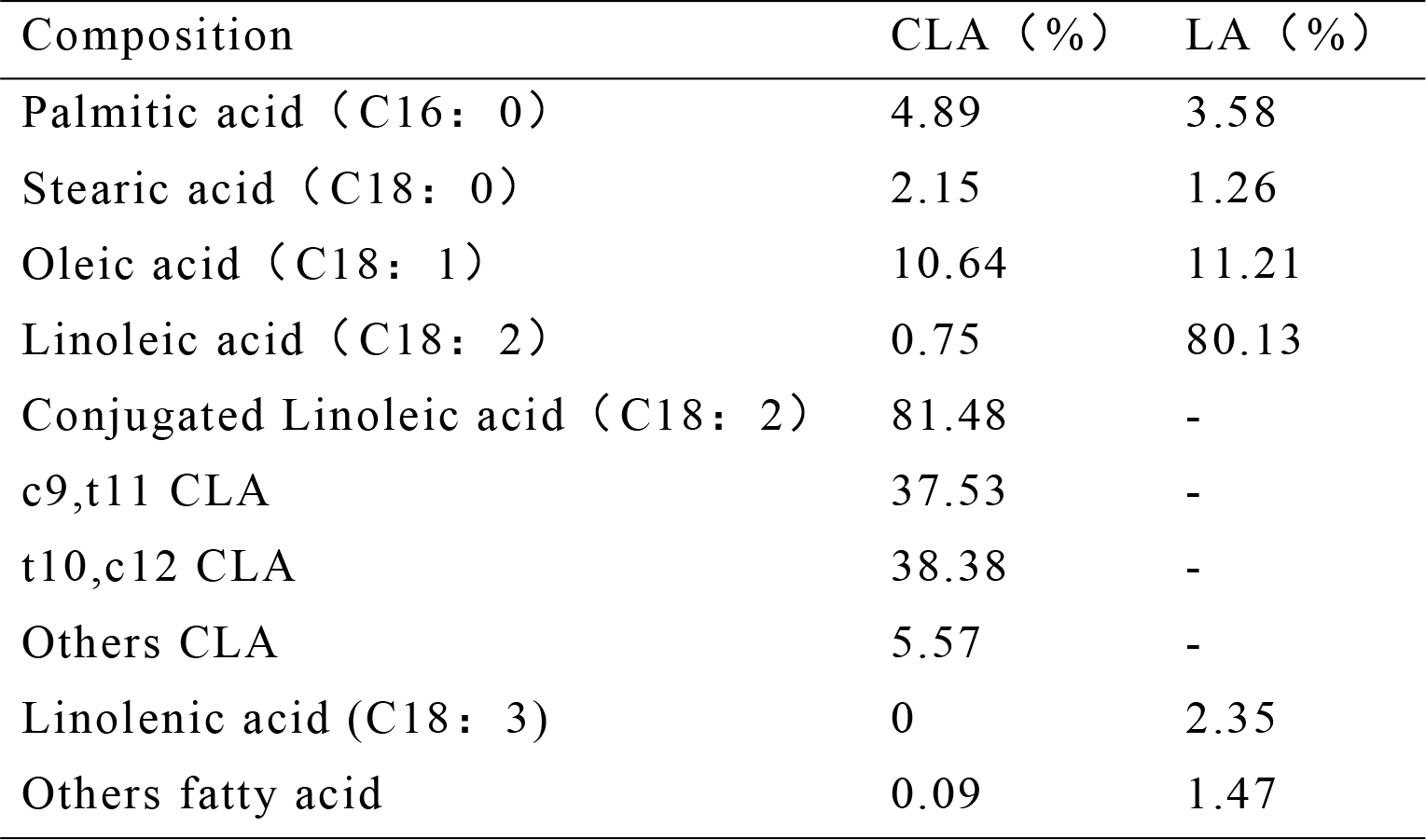
composition of LA oil and CLA oil

| Composition | CLA (%) | LA (%) |
| --- | --- | --- |
| Palmitic acid (C16: 0) | 4.89 | 3.58 |
| Stearic acid (C18: 0) | 2.15 | 1.26 |
| Oleic acid (C18: 1) | 10.64 | 11.21 |
| Linoleic acid (C18: 2) | 0.75 | 80.13 |
| Conjugated Linoleic acid (C18: 2) | 81.48 | - |
| c9,t11 CLA | 37.53 | - |
| t10,c12 CLA | 38.38 | - |
| Others CLA | 5.57 | - |
| Linolenic acid (C18: 3) | 0 | 2.35 |
| Others fatty acid | 0.09 | 1.47 |

In experiment one, all dams were fed the laboratory chow diet for the first 3 days of lactation. On day 4 postpartum, six lactating mice were randomly assigned to the control group (CON group, 3.0% LA oil), the low level CLA group (L-CLA group, 1.0% CLA + 2.0% LA), the median level CLA group (M-CLA group, 2.0% CLA + 1.0% LA), and the high level CLA group (H-CLA group, 3.0% CLA) from day 4 to 10 postpartum. The litter number was normalised to eight at postpartum. Dam weight, food intake, and the eight pups weight were determined daily between days 4 to 10 postpartum. The accurate assessment of food intake was achieved by accurately collecting spilled food from cages. Based on the previous day’s intake, a sufficient amount of feed was offered to ensure at least 10% refusal. On day 10 postpartum, pups were separated from dams for 2 h before feeding to ensure maximal accumulation of milk in the mammary gland. Subsequently, all dams were milked by suction. Pups were killed by cervical dislocation, and their abdomens were opened and milk clots within the stomach were weighed and stored at −78°C until analysis [12,23]. The milk lipid concentration was measured gravimetrically after chloroform-methanol extraction by a modified Folch method [24]. Protein concentration was determined by the Biuret method with casein as a standard. The milk lactose concentration was estimated by Cocciardi method [25]. Dams were killed on day 10 of lactation. Mammary and liver tissue from lactating mice was excised and weighed, frozen in liquid nitrogen, and stored at −80°C. Triglycerides in liver were extracted and analysed using a Tissue Triglyceride Assay Kit (E1013, Applygen Technologies Inc) according to the manufacturer’s protocol.

In experiment two, dams received 3.0% LA oil (CON group) or 3.0% CLA oil (H-CLA group) from day 4 to day 10 of lactation. Feeding conditions were the same as in the first experiment. Blood was collected from dams and analysed in the First Affiliated Hospital of Zhengzhou University. Plasma insulin concentrations were quantified using a Mouse Insulin Ultra-sensitive ELISA Kit (BC1710, Solarbio). For ITTs, on the 11th day of lactation, all pads were renewed, and dams were starved for 6 h and weighed. Each dam was intraperitoneally injected with insulin (Novolin) at a dose of 0.75 U/kg body weight. Before injection, the baseline blood glucose concentration was measured and recorded as 0 min. After injection of insulin, the blood glucose level in the tail vein was measured at 15, 30, 60, 90 and 120 min using a TIDEX glucose analyser (Sankyo, Tokyo). The rate constant for each ITT (KITT) was calculated using the equation KITT (%/min) = 0.693 / t(1/2), where t(1/2) was calculated from the slope of plasma glucose concentration during 15–90 min after administration of intravenous insulin.

### Statistical analysis

Results are reported as mean ± standard deviation (SD). Data statistics and differences among groups were analysed by SPSS 13.0 (IBM, USA). All experiments were conducted in triplicate and repeated at least twice. Data were analysed for significant differences using a general linear model of repeated measures, one-way analysis of variance (ANOVA), or Student’s t-tests. A *p*-value <0.05 was considered statistically significant.

## Results

### Dam body weight and food intake

Dam body weight increased progressively, and revealed a linear effect over time (*p* <0.01, Fig. 1A), but there were no significant differences between different treatment groups. Significant effects of treatment and time were observed for food intake of dams (*p* <0.05, Fig. 1B). Within the CLA treatment groups, there was a significant difference in food intake between the L-CLA and M-CLA groups, and the H-CLA group.

**Figure 1.**
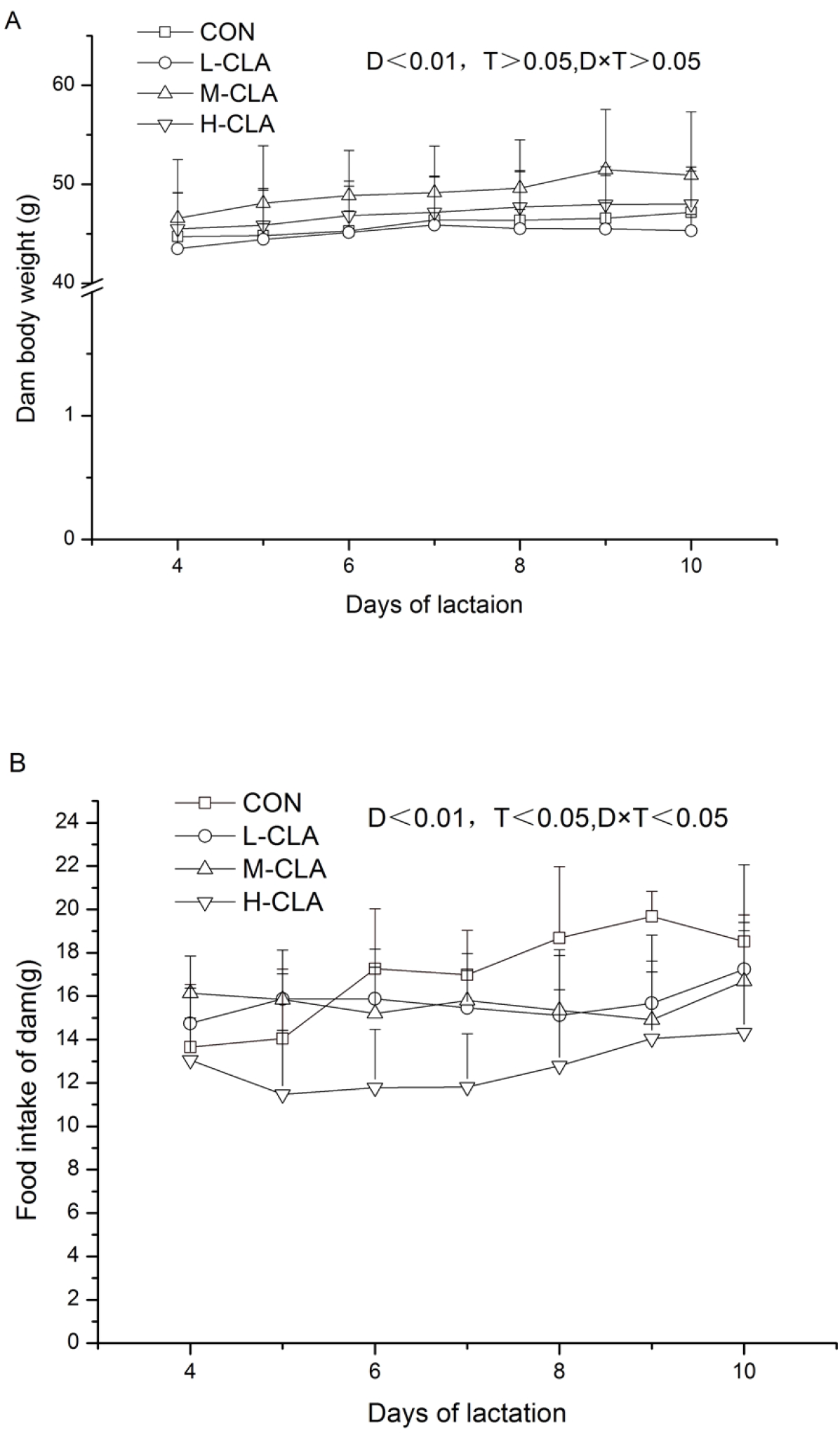
Daily body weight and food intake of dams fed different doses of CLA. Values are means ± SD (n = 6). (A) Dam body weight during day 4 to day 10 of lactation. (B) Food intake during day 4 to day 10 of lactation. CON group, 342.0% LA oil; L-CLA group, 1.0% CLA + 2.0% LA; M-CLA group, 2.0% CLA + 1.0% LA; H-CLA group, 3.0% CLA; T, effect of CLA treatment. D×T, time and treatment interaction.

### Pup body weight

The eight pups weight was measured daily and the results are shown in Fig. 2. Significant effects for both both time and treatment were observed for pup weight gain over the feeding period. Although the pup weight in all four groups increased from day 4 to day 10, pups in the H-CLA group grew more slowly, and at the end of the study, the eight pup weight of the H-CLA group was significantly lower than that of the other groups (39.38 vs. 54.47, 54.95, and 51.13, *p* <0.01) by day 10 of lactation.

**Figure 2.**
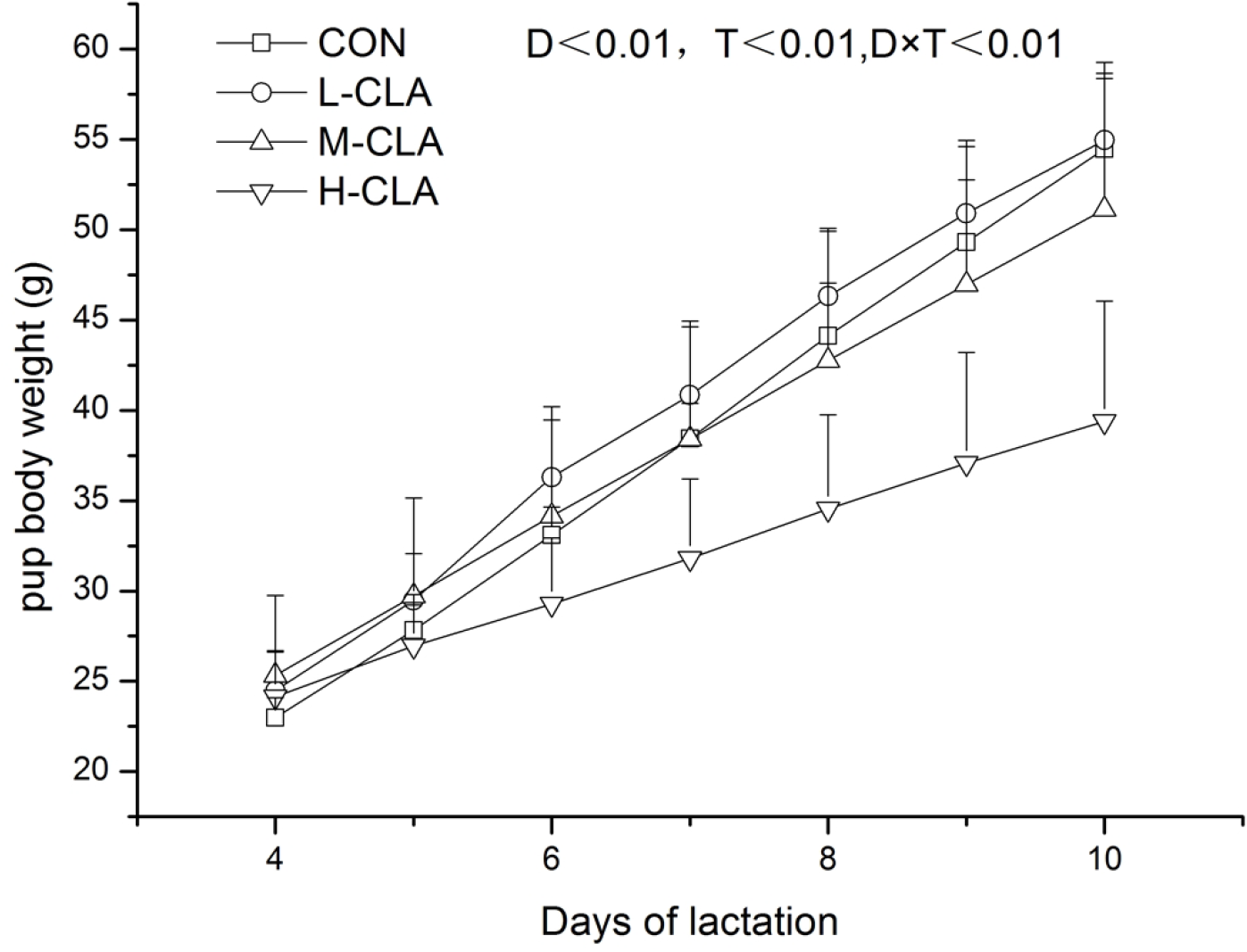
Daily weight of nursing pups fed different doses of CLA during day 4 to day 10 of lactation. Values are means ± SD (n = 6). CON group, 342.0% LA oil; L-CLA group, 1.0% CLA + 2.0% LA; M-CLA group, 2.0% CLA + 1.0% LA; H-CLA group, 3.0% CLA; D, effect of lactation time, T, effect of CLA treatment; D×T, time and treatment interaction.

### Milk and liver

The milk clot weight obtained from the stomach ranged between 0.91 g and 0.56 g/suckling pup. The milk fat content of the H-CLA group was significantly lower than that of the other groups. There were no significant differences in protein and lactose concentrations in milk between CLA and control groups (Table 2). Similarly, there were no differences in the weight of mammary tissue between these groups. Liver weight was increased gradually with increasing dose of CLA, and there was a significantly higher TG content in the liver of H-CLA group mice compared with other groups (Table 2).

**Table 2.** Effect of CLA on the milk and liver of dams

|  | CON group<sup>#</sup> | L-CLA group | M-CLA group | H-CLA group |
| --- | --- | --- | --- | --- |
| Milk clotweight (g/pup ) | 0.89±0.27<sup>a</sup> | 0.91±0.12<sup>a</sup> | 0.85±0.23<sup>a</sup> | 0.56±0.29<sup>b</sup> |
| Milk fat (mg/100mg) | 15.42±1.43<sup>a</sup> | 14.16±1.96<sup>a</sup> | 13.37±2.30<sup>a</sup> | 11.88±1.99<sup>b</sup> |
| Milk protein (mg/100mg) | 18.79±1.35 | 20.19±2.30 | 17.96±3.71 | 19.87±3.32 |
| Milk lactose (mg/100mg) | 2.03±0.56 | 1.89±1.45 | 2.26±0.85 | 1.93±1.30 |
| Mammary Weights (g) | 3.88±0.69 | 4.21±0.79 | 4.39±1.23 | 3.96±1.06 |
| liver Weights (g) | 2.91±0.29<sup>a</sup> | 3.02±0.41<sup>a</sup> | 3.63±0.55<sup>ab</sup> | 4.65±0.59<sup>b</sup> |
| TG of liver (mg/g tissue) | 46.41±27.07<sup>a</sup> | 94.51±32.78<sup>ab</sup> | 92.5±34.16<sup>ab</sup> | 230.14±80.54<sup>b</sup> |

<sup>a,b</sup>Values followed by different letters in the same row indicate significant differences
( $p < 0.05$ ). Values are means ± SD.

### Blood metabolite analysis

Similar to experiment one, in experiment two, the weight of pups in the high dose CLA group was significantly lower than that in the control group (Fig. 3). CHO, HDL and LDL levels in serum were unaffected, but plasma T and γ -GT activity in the H-CLA treatment group were higher than in the control group (*p* <0.05), and glucose was also increased (*p* = 0.07, Table 3). Insulin activity was significantly higher in the H-CLA group. Interestingly, CLA feeding resulted in higher RBC and HGB values in the blood of mice in the H-CLA group.

**Table 3.**
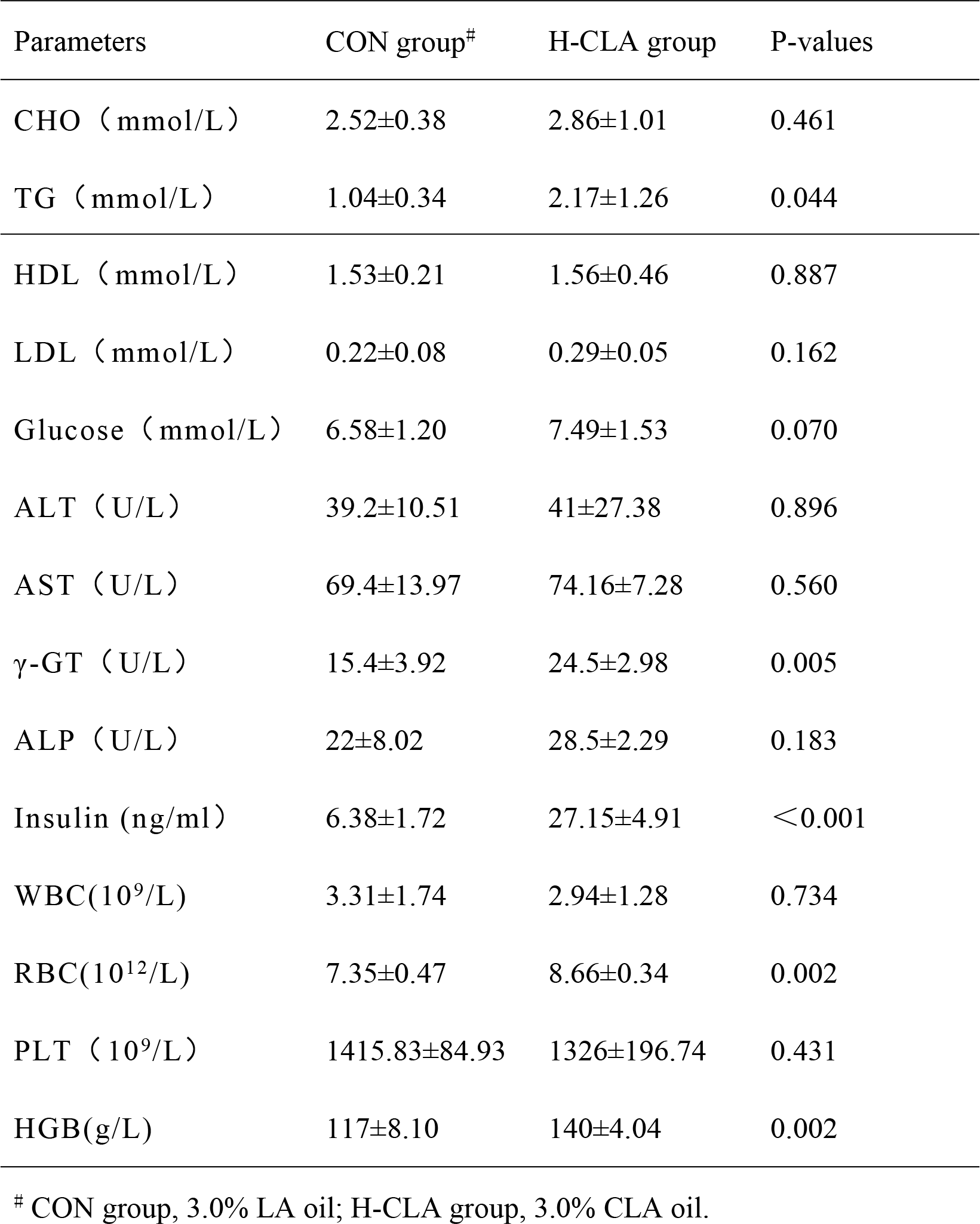
Effect of a high dose of CLA on the blood of dams

**Figure 3.**
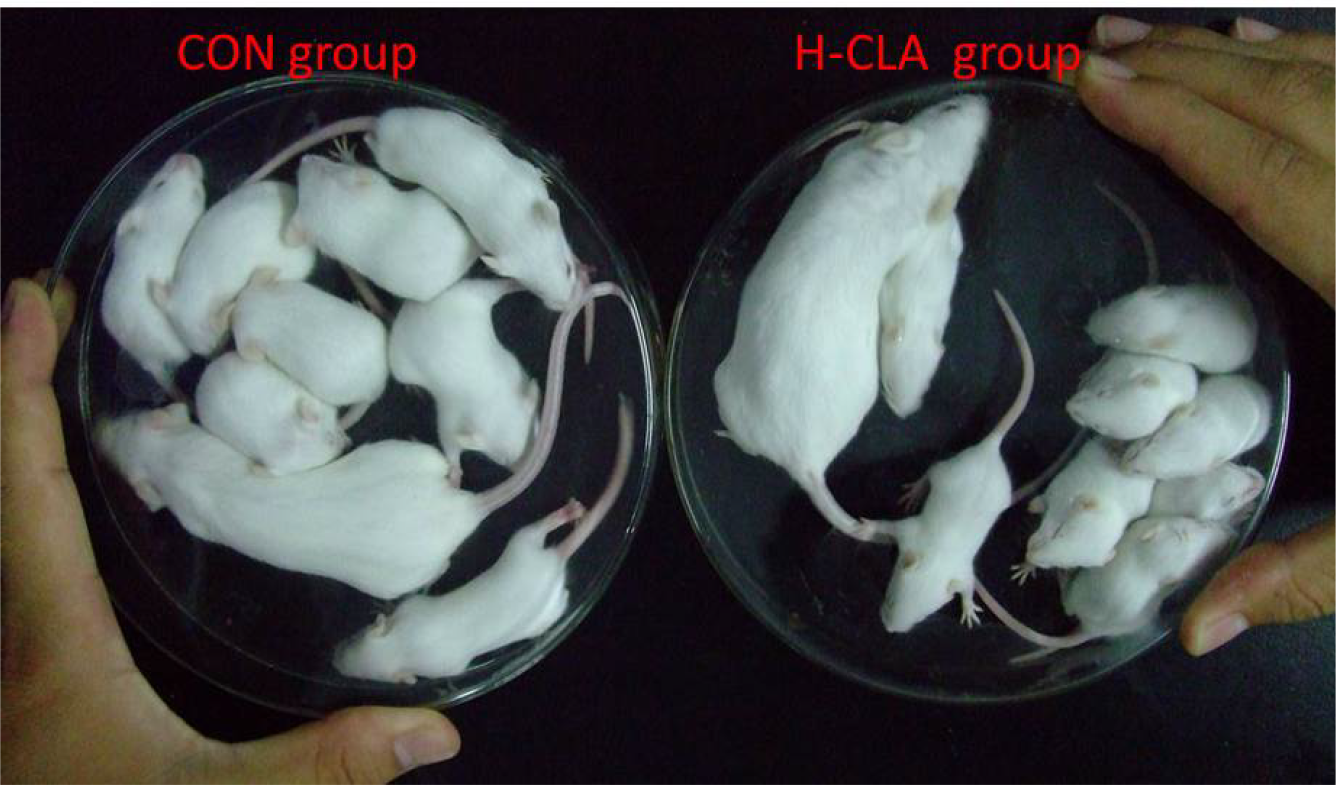
Dams and eight pups of CON and H-CLA groups in experiment two at day 10 of lactation.

**Figure 4.**
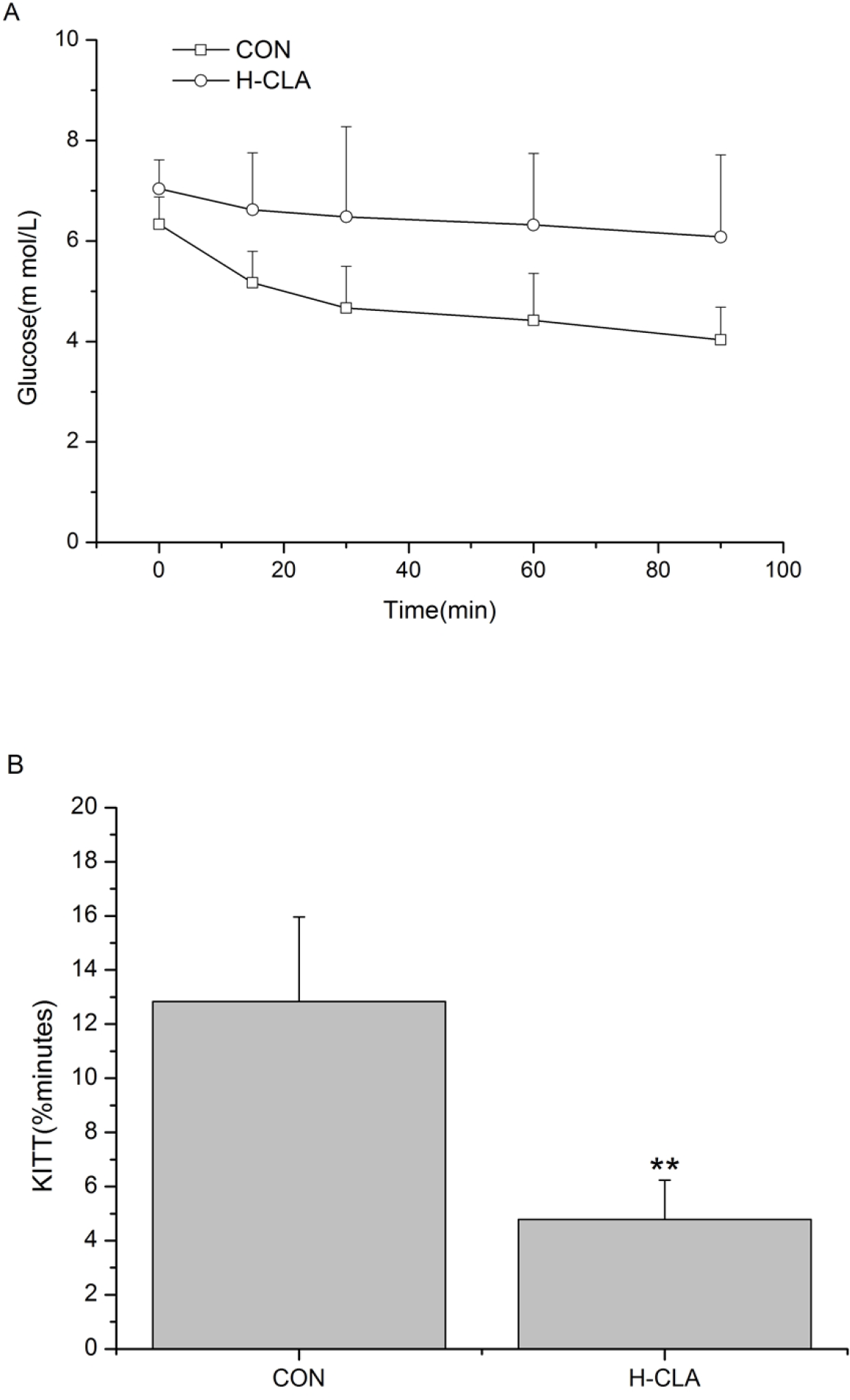
Insulin tolerance tests (ITTs) of lactating mice in CON and H-CLA groups. Values are means ± SD (n = 6). (A) ITTs of lactating mice. Each dam was injected with insulin at a dose of 0.75 U/kg body weight. Blood glucose level was measured at 0, 15, 30, 60, 90 and 120 min. (B) Calculation of KITT (%min) after a dose of 0.75 U of insulin per kg of body weight. CON group, 2.0% LA oil; H-CLA group, 2.0% CLA; \*\**p* <0.01.

### Insulin tolerance testing

At 0 min, the blood glucose level in the control group was 6.62 mmol/L, and that of the CLA group was 7.04 mmol/L (Fig. 3). After injection of insulin, glucose in the control group decreased to 4.03 mmol/L at 90 min, while glucose levels in the H-CLA group were not obviously decreased (6.08 mmol/L). As shown by the decrease in KITT, feeding mice with an high dose of CLA resulted in a significant insulin resistance state.

## Discussion

CLA can reduce fat deposition, weight gain, and food/energy intake in mice [26]. In our present study, CLA treatment decreased food intake but did not reduce dam weight. Although the mechanism is not clear, it is likely related to the lactation stage, since mammary glands are the most active lipid-secreting organ in lactating mice [27]. Chin (1994) reported that increased maternal intake of CLA during lactation enhanced the growth rate of suckling neonatal rats [28]. In contrast to the stimulatory effect of CLA, mixtures of CLA [19] or isoforms [18] caused litter pups to grow slowly or even die due to decreased milk fat. These contradictory results might be due to differences in the content or isomeric distribution of the CLA supplement. In our present study, 2% CLA diet fed to KM mice during lactation decreased both milk weight and milk fat content. Milk is the only source of nutrients for suckling pups; hence it was not surprising that a reduction in the fat and weight of milk were accompanied by reduced pup weight during lactation. These results are consistent with those of other studies on mice [12]. These findings confirmed that a high dose of CLA lowered the content of milk fat and pup growth rate.

CLA has been reported to generate fatty liver, which could be a consequence of increased lipogenesis in the liver to compensate the reduction in fat deposition in adipose tissue [26]. We assessed the liver of dams and observed increased liver weight and TG content in mice given a high dose of CLA. Kadegowda (2010) reported that mice treated with CLA displayed decreased milk fat and increased liver weight due to hepatic steatosis [29], similar to our current results. The effects of CLA supplementation on mammary fat synthesis have received a great deal of attention [30,31], but the potentially adverse effects of CLA on the liver and lipid metabolism have been largely ignored in lactating animals. We suggest that severe lipid accumulation occurs in the liver when lactating mice are given a high dose of CLA. This was also confirmed by increased TG content and γ-glutamyl-transferase (γ-GT) activity in the serum of the high CLA group.

There have been many reports regarding CLA causing insulin resistance [7]. In the present study, ITTs clearly indicated that a high dose of CLA fed to mice caused insulin resistance. In lactating mice, glucose is the main precursor for the synthesis of lactose and triglyceride substrates. During the lactation stage, mammary glands have increased demand for glucose, and are highly sensitive to insulin, while other organs are relatively insensitive to insulin [32]. In this study, lactating mice fed with CLA displayed aggravated insensitivity to insulin and exhibited insulin resistance, implicating insulin as a key factor in CLA-mediated regulation of milk fat and lactation.

Interesting, we observed for the first time that a high dose of CLA can result in higher RBC and HGB levels in the blood of lactating mice. RBC and HGB are both positively correlated with insulin resistance and NAFLD [33]. Thus, in the present study, higher RBC and HGB may be a consequence of insulin resistance and fatty liver in mice receiving an high dose of CLA.

In summary, a high level of CLA ingestion during lactation decreases milk fat and increases lipid content in the liver. Therefore, CLA is a dietary factor with a propensity to affect hepatic metabolism during lactation. Little is known about the metabolic changes in animals during lactation, especially the possibility of a very complex regulatory mechanism between lipid metabolism in the liver and mammary lactation. The present findings provide a platform for further studies on the exact mechanism and the safe use of CLA.

